# Functional maps of a genomic locus reveal confinement of an enhancer by its target gene

**DOI:** 10.1101/2024.08.26.609360

**Authors:** Mathias Eder, Christina J.I. Moene, Lise Dauban, Christ Leemans, Bas van Steensel

**Author notes:** These authors contributed equally to this work (Mathias Eder, Christina J.I. Moene).

## Abstract

Genes are often activated by enhancers located at large genomic distances. The importance of this positioning is poorly understood. By relocating promoter-reporter constructs into >1,000 alternative positions within a single locus, we dissected the positional relationship between the mouse *Sox2* gene and its distal enhancer. This revealed an intricate, sharply confined activation landscape, in which the native *Sox2* gene occupies an optimal position for its activation. Deletion of the gene relaxes this confinement and broadly increases reporter activity. Surprisingly, the confining effect of the *Sox2* gene is partially conferred by its ∼1 kb coding region. Our local relocation approach provides high-resolution functional maps of a genomic locus and reveals that a gene can strongly constrain the realm of influence of its enhancer.

## MAIN TEXT

Mammalian enhancers can activate genes across large genomic distances, up to hundreds of kilobases (*1, 2*), presumably by contacting their target promoter through folding of the chromatin fiber. Indeed, enhancers tend to have increased contact with the promoters they regulate (*3*), but it is unclear whether this is a cause or effect of their function. In addition, enhancer – promoter pairs often reside within the same topologically associated domain (TAD), a region of preferential self-interaction in the genome that is often demarcated by CTCF binding sites (*4, 5*). These TADs are thought to form functional domains (*6, 7*), but it is largely unclear whether enhancers can activate promoters anywhere within a TAD, or whether some positions within a TAD are more favorable than others. Recent studies in two loci with little three-dimensional (3D) structure indicated that promoter activity increases with decreasing genomic distance to its enhancer (*8, 9*). However, in more structured genomic loci the effect of position on enhancer – promoter communication has remained elusive. Here, we addressed these questions by constructing high-resolution maps of promoter activity as function of position in a native locus with pronounced 3D folding.

### High throughput relocation by Sleeping Beauty transposition

To unravel the logic that underlies the linear arrangement of regulatory elements, it is necessary to alter the genomic position of such elements systematically and probe the functional consequences. Here we provide such a systematic transplantation of regulatory elements by adapting a methodology based on the cut-and-paste mechanism of transposable elements (*9–11*). When mobilized from the genome by its transposase, the Sleeping Beauty (SB) transposon excises and re-integrates, usually within ∼1-2 Mb from the original location (**Fig. 1A**) (*10, 12*). Thus, any sequence inserted between the two SB inverted terminal repeats (ITRs) can be randomly relocated (“hopped”) throughout a genomic locus. We developed a workflow to generate and map thousands of integrations in a single locus, and link the locations to the expression of a transcribed reporter, yielding high-resolution functional maps of the local regulatory landscape.

**Fig. 1.**
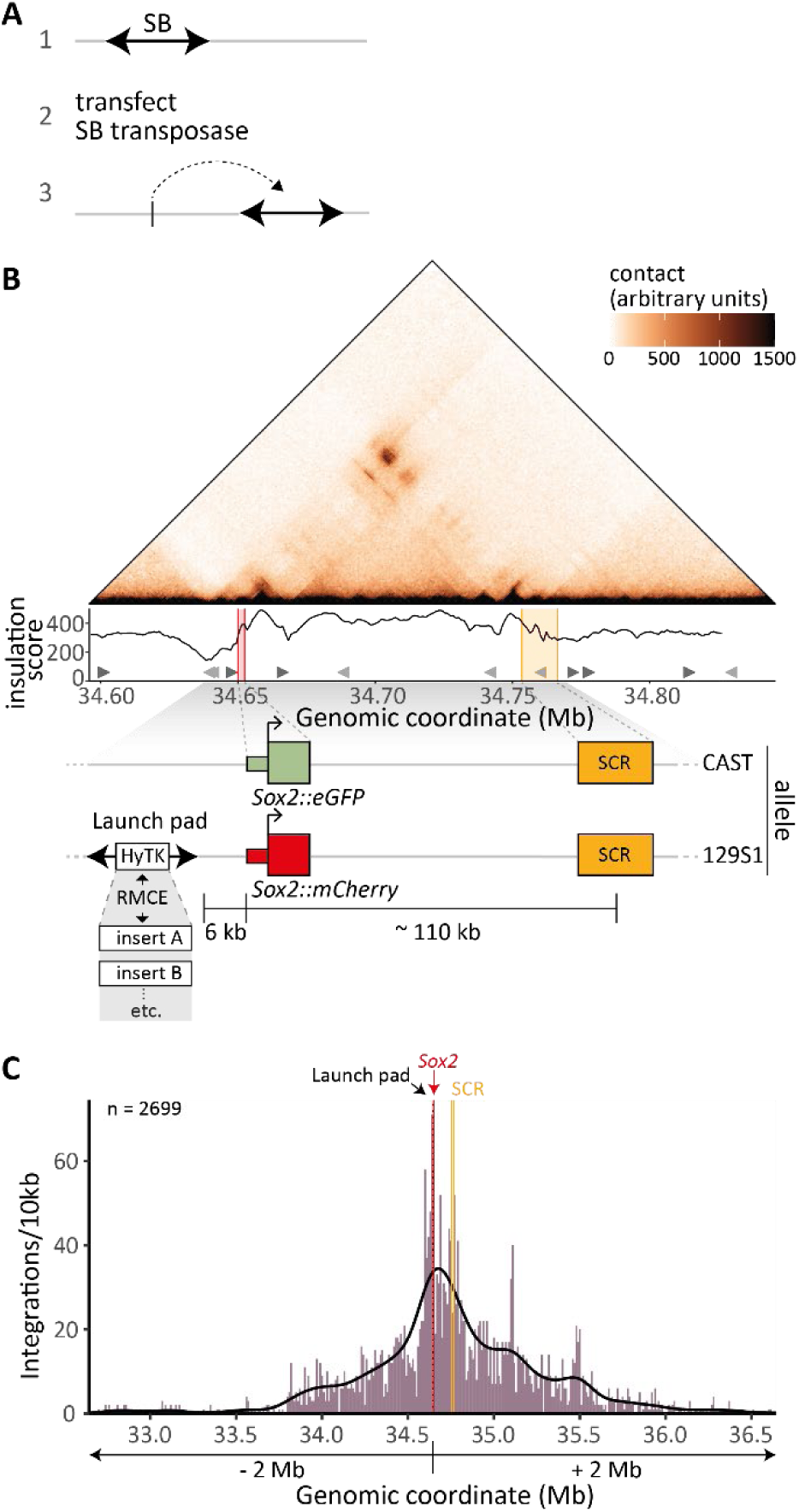
Sleeping Beauty mobilization creates integrations covering the *Sox2* locus. **(A)** Cut-and-paste local relocation of a Sleeping Beauty (SB) transposon by transient expression of SB transposase. **(B)** Overview of the *Sox2* locus. Top to bottom: contact map in mESCs (region-capture micro-C data [(*18*)], 1000-bp resolution) and insulation scores based on the same data; positions and orientations of CTCF binding sites (triangles); and positions of SCR and eGFP/mCherry tagged *Sox2* alleles, the SCR, and a launch pad (129S1 allele) containing SB with a recombination cassette. **(C)** Distribution of hopped SB carrying a 282 bp inert DNA sequence, mobilized from the launch pad; n, total number of integrations mapped in the plotted region.

For this study we chose the *Sox2* locus in mouse embryonic stem cells (mESCs). In these cells, *Sox2* expression is primarily determined by the *Sox2* control region (SCR), which is a cluster of enhancers located 110 kb downstream of the gene in the same TAD (*13–17*). Within this TAD, the *Sox2* gene has been found to preferentially contact the SCR (*18*) (**Fig. 1B**). The 1.5 Mb region around *Sox2* contains no other protein-coding genes (*13*). To ease genetic engineering, we used F1-hybrid (*129/Sv:CAST/EiJ*) mESCs, in which the *CAST* and the *129S1* alleles can be distinguished by a high density of sequence polymorphisms. We tagged both alleles of *Sox2* with either *eGFP* (*CAST*) or *mCherry* (*129S1*) to be able to quantify allelic *Sox2* expression by flow cytometry (**Fig. S1A**). To confirm the regulation by the SCR, we transfected the dual-tagged cell line with Cas9 and two gRNAs that cause deletion of the SCR. In line with the reported key role of the SCR (*13, 14, 16*), a proportion of cells lost more than 90% of *eGFP* or *mCherry* expression, most likely corresponding to a deletion of the SCR on the corresponding allele (**Fig. S1B**).

Next, we used Cas9 editing to create a “launch pad” by integrating a SB transposon cassette 6 kb upstream of the *Sox2::mCherry* allele (−116 kb relative to the SCR, **Fig. 1B**). The cassette contained a double selection marker (mPGK-HyTK) flanked by a pair of heterotypic Flp-recombinase recognition sites (FRT/F3), enabling us to easily change the DNA sequence within the transposon by recombination-mediated cassette exchange (RMCE) (**Fig. S1C**).

In a proof-of-principle experiment, we induced hopping of the SB cassette containing an arbitrary 282 bp sequence (see methods). After culturing the cells for one week to dilute the transposase-containing plasmid and prevent further re-hopping events, we isolated and expanded 200,000 cells and mapped the integrations using a Tn5-based sequencing approach (*19*). In some of the cells, the transposon was never mobilized, which is reflected by 33% of the mapping reads originating from the launch site. Of all the hopped integrations, 72% were located on the same chromosome as the *Sox2* locus (chr3) (**Fig. S1D**). Allele-specific mapping indicated that most integrations happened in *cis* (**Fig. S1E**). Within −/+ 2 Mb of the launch pad we mapped 2699 unique integrations (**Fig. 1C**), which were equally distributed between the forward (1342) and reverse (1357) orientations (**Fig. S1F**). We note that this mapping is not exhaustive and the actual pool likely contains more integrations, since more than 70% of the integrations were only supported by one unique mapping read (**Fig S1G**). This experiment demonstrates the effectiveness of SB hopping to generate thousands of genomic integrations.

### A highly detailed activation landscape of the *Sox2* locus

Next, we set out to generate a detailed map of the expression potential of the *Sox2* locus, *i.e.*, to determine quantitatively how the location throughout the locus affects the ability of the *Sox2* promoter to be activated. For this we engineered a reporter comprising 1.9 kb of the *Sox2* promoter (excluding a promoter-proximal CTCF binding site) driving the expression of mTurquoise2 (*mTurq*), a blue fluorescent protein (**Fig. 2A**). We integrated this reporter into the launch pad using RMCE. This resulted in low but detectable reporter expression, without affecting endogenous *Sox2::mCherry* expression (**Fig. S2A**, reporter −116 kb).

**Fig. 2.**
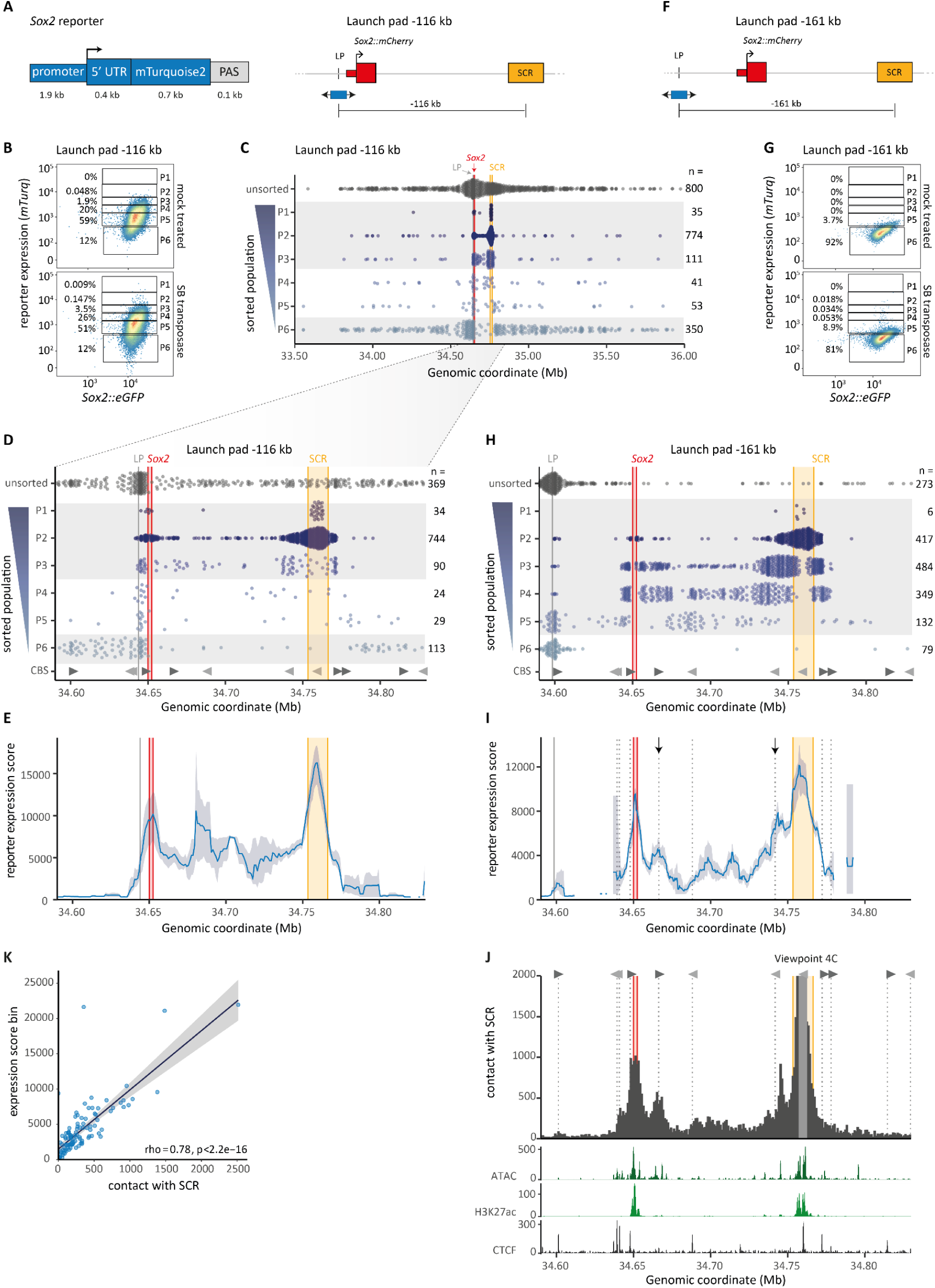
A highly detailed activation landscape of the Sox2 locus. **(A)** Left panel: design of mTurquoise2 reporter driven by *Sox2* promoter and 5’UTR, followed by SV40 polyadenylation signal (PAS). Right panel: this reporter was inserted in the −116 kb launch pad (LP), upstream of the mCherry-tagged endogenous *Sox2* allele. **(B)** Expression levels of reporter (mTurq) and *Sox2::eGFP* (control) in mock-transfected cells and in cells transfected with SB transposase, measured by FACS. Boxes mark sorting gates used to obtain cell populations P1-P6 in **C-D**. The percentage of cells in each gate is indicated. Representative result of three biological replicates. **(C)** Mapped integrations from a large pool of unsorted cells (ctrl) and the six sorted populations (P1-P6), mobilized from the −116 kb launch pad. Each dot indicates one mapped integration, number of plotted integrations in each population is indicated on the right. Combined data from three biological replicates. Grey shading highlights populations that are most different from the starting expression of the unhopped reporter. LP, launch pad position. **(D)** Zoom-in of C. Grey triangles indicate CTCF binding sites (CBS) and their orientation. **(E)** Expression score derived from data shown in **D**, smoothened using a 10 kb running window shifted in 1 kb steps and only plotted when the window contains 3 or more SB integrations. Shaded region indicates 95% confidence interval. **(F-I)** Same as **A-B, D-E**, except that the reporter was mobilized from a launch pad at position −161 kb. Data are from 2 biological replicates combined. Expression scores in **I** are calculated using a 5 kb running window shifted in 500 bp steps. Arrows mark two peaks near CBSs. **(J)** Annotation of the locus. Top: virtual 4C contact profile (from RCMC data, (*18*)) with the center of the SCR (indicated in lightgrey) as viewpoint, at 1 kb resolution. Dashed lines indicate CBSs and grey triangles indicate their orientation. Bottom: ATAC-seq signal (*20*), H3K27ac ChIP-seq p-value signal (*21*) and CTCF ChIP-seq coverage (own data), smoothened using a 500bp running window. **(K)** Correlation between expression score and virtual 4C contact with the core SCR, for the datasets from −161 kb launch pad, per 1 kb bin on the genomic region plotted in **H-J**. Only bins with 3 or more SB integrations are included. Rho is Spearman’s correlation coefficient.

We then induced SB hopping and monitored reporter fluorescence by flow cytometry. Compared to the non-hopped control, the resulting cell pool showed a broadened distribution of reporter fluorescence, indicating that the relocation at least in part affected reporter expression (**Fig. 2B**). We then split the full range of reporter expression values into six gates (P1-P6) and sorted pools of cells from each gate (**Fig. 2B**), and subsequently mapped the locations of the reporters in each pool. As a control, we also mapped the locations in unsorted cells. Since the integration patterns were reproducible between biological replicates (**Fig. S2B-C**), we combined the replicates for further downstream analysis.

As we observed previously with the arbitrary insert sequence (**Fig. 1C**), reporter integrations in the control pool are mostly concentrated in a 2 Mb region surrounding the launch site (**Fig. 2C**, top track). In contrast, the cell pools with different expression levels yielded clearly distinct distributions (**Fig. 2C-D**, P1 – P6 tracks). Reporters with the highest activity (P1) were concentrated almost exclusively in the center of the SCR or close to the endogenous *Sox2* gene, while the second-most active reporters (P2) were more dispersed around these regions. Intermediate activity (P3 – P5) occurred predominantly near the launch site and between *Sox2* and SCR. Finally, silent (P6) reporters were almost exclusively located outside the *Sox2*-SCR TAD.

We converted these raw data into a reporter expression score, which is the average expression level estimated in a sliding window across the locus (**Fig. 2E**, see Methods). This graph highlights that the SCR and the region immediately surrounding the endogenous *Sox2* gene are the most optimal locations for transcriptional activity. Precise fluorescence measurements in a panel of clonal cell lines with 36 unique integrations confirms this peak in activity at the center of the SCR and validates the pattern of expression scores inferred from the sorted pools (**Fig. S2D**). Reporter integrations in the center of the SCR display a 10-fold upregulation in reporter expression compared to the −116 kb launch pad position. A remarkable lack of reporter expression occurs outside of the *Sox2*-SCR TAD, with a very sharp drop-off (within about 10 kb) on either side (**Fig. 2E**). A plateau of intermediate expression is found between *Sox2* and SCR. Together, this detailed landscape points to a strong confinement of the realm over which the SCR is able to activate transcription.

The relatively large confidence intervals of the expression scores limited interpretation of the fine-scale pattern between *Sox2* and SCR (**Fig. 2E**). This is due to a relative paucity of reporter integrations in this range. The distribution of reporter expression in cells where the reporter was not mobilized, overlaps the medium-level expression gates (P4 – P5, **Fig. 2C**), leading to a low number of sorted integrations for those gates. To overcome this, we repeated the hopping experiments starting from a launch pad 51 kb upstream of *Sox2::mCherry* (−161 kb relative to the SCR) (**Fig. 2F**). At this position the reporter showed no measurable expression (**Fig. S2A**), and hence from this launch pad it should be easier to detect hopping events in the medium-level expression bins. Indeed, this was the case (**Fig. 2G-H**). The resulting expression score map (**Fig. 2I**) has narrow confidence intervals within the *Sox2*-SCR TAD (at a resolution of 5 kb), at the cost of a paucity of data outside this range.

This high-resolution map confirmed that the highest reporter expression occurs at the SCR and around the endogenous *Sox2* gene (**Fig. 2I**). Reporter activity throughout the region is independent of the orientation of the reporter (**Fig. S2E**). Within the *Sox2*-SCR TAD, the map reveals an intricate activity landscape with several local peaks of reporter expression, two of which overlap CTCF binding sites (**Fig. 2I**, black arrows). Strikingly, this landscape closely matches a map of SCR contact frequencies that was obtained in the absence of reporter integrations (*18*) (**Fig. 2J**, top panel; **Fig. 2K**). Maps of open chromatin (ATAC-seq (*20*)) and the histone mark H3K27ac (*21*), both thought to be features of regulatory elements (*22*), showed only partial overlap with the fine pattern of reporter activity (**Fig. 2J**, bottom panel). These results indicate that reporter expression primarily follows contact frequency with the SCR, suggesting that these contacts are a major determinant of quantitative expression levels.

### Random deletion of the endogenous *Sox2* gene boosts reporter expression

Close examination of the flow cytometry data of the transposase-transfected reporter cells (−116 kb launch pad) revealed a small but distinct population of cells with high reporter expression but no *Sox2*::*mCherry* expression (**Fig. 3A**). We isolated a panel of clones with these unusual characteristics and mapped the SB location. Surprisingly, in these clones the left ITRs were still located at the launch site (upstream of *Sox2::mCherry*), while the right ITRs were linked to sequences distributed throughout the *Sox2* locus (**Fig. 3B, S3A**). This suggests that the genomic sequence between the launch site and integration site has been lost. Sanger sequencing confirmed the loss of the 129S1 allele in this region (**Fig. S3B**). We hypothesize that in rare instances, errors in the SB excision-reinsertion process can cause such a deletion, which can lead to the loss of the *Sox2::mCherry* gene.

**Fig. 3.**
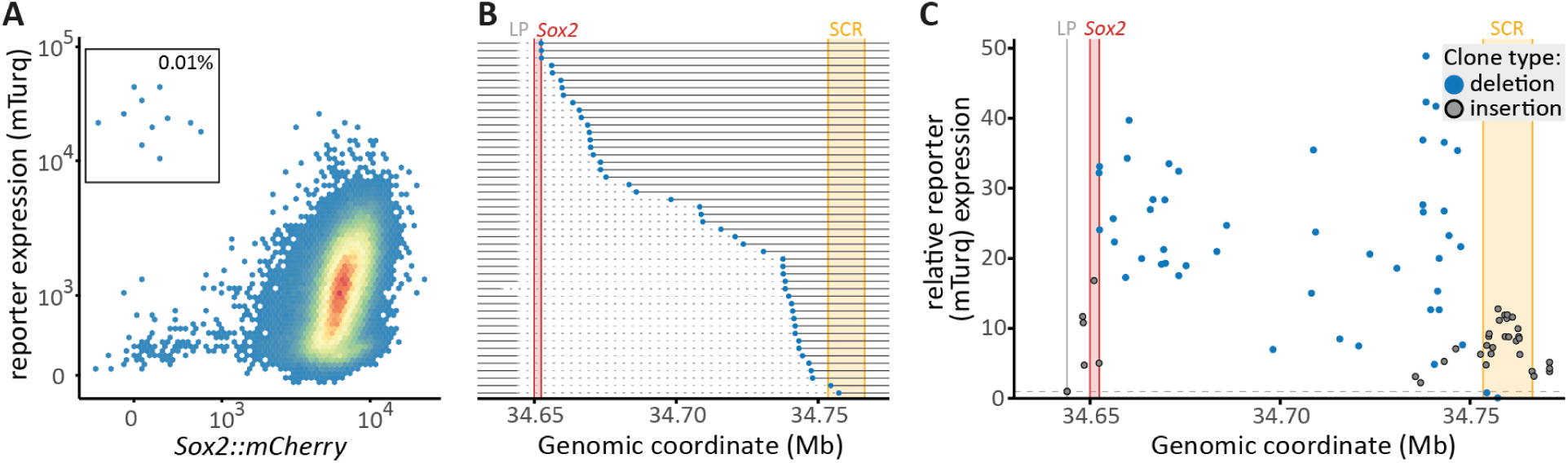
Hopping-induced deletions of the endogenous Sox2 gene increase reporter expression. **(A)** Expression of *Sox2::mCherry* and *Sox2* reporter (mTurq) after mobilization of SB from the −116 kb launch pad, first biological replicate (also shown in **Fig. 2B**). Box indicates rare *Sox2::mCherry*-negative cells with high reporter expression. **(B)** Unique deletions inferred from mapping in *Sox2::mCherry*-negative sorted clones (**Fig. S3A**), sorted by deletion size. Horizontal dashed lines indicate individual deletions. Dots indicate the locations of the re-insertions. Almost all deletions start at the launch pad position. **(C)** Relative reporter expression of clones with a hopping-induced deletion (blue dots, location corresponds to the end of the deletion) or regular insertion (grey, same data as in **Fig. S2D**, bottom).

Strikingly, clones with such hopping-induced deletions exhibited very high reporter expression, up to 49-fold higher than the −116 kb reporter clone (**Fig. 3C**). Reporters close to the SCR (within 20 kb) were expressed about 5-fold higher when a deletion was present, compared to regular insertions in the same area. Even the smallest deletions of 8.5 kb, removing only the endogenous *Sox2::mCherry* gene, resulted in dramatically increased reporter expression. These data suggest that the endogenous *Sox2* gene strongly limits reporter expression when present. Other sequences within the *Sox2*-SCR range may further modulate this effect, indicated by the variable increases in expression across the deletions, which show no correlation with distance to the SCR (Spearman’s rho = 0.19, p = 0.19). In summary, these fortuitous rare deletions suggested that the endogenous *Sox2* gene strongly reduces activity of the reporter gene throughout the *Sox2*-SCR range.

### *Sox2* gene strongly controls the regulatory landscape of the enhancer

To verify this interpretation, we used CRISPR/Cas9 to delete the entire *Sox2* gene, including its promoter. Furthermore, to test whether the CTCF binding site (CBS) just upstream of the *Sox2* promoter (which was also lost in the hopping-induced deletions) might play a role, we deleted either the *Sox2* promoter and gene body (Δ*Sox2*), or the same region including this CBS (ΔCBS-*Sox2*). The guide RNAs used for these deletions were not allele-specific, and thus could independently delete either the *Sox2::eGFP* or *Sox2::mCherry* allele (homozygous *Sox2* deletion causes mESC differentiation (*23*)). By FACS analysis we identified cells with either heterozygous deletion and measured the respective impacts on mTurq reporter activity (**Fig. 4A)**. We performed these experiments in three cell lines, with the reporter located at positions −161 kb (no detectable activity), −116 kb (low activity), or −39 kb (between *Sox2* and SCR, medium activity) (**Fig. S4A**) relative to the SCR.

**Fig. 4.**
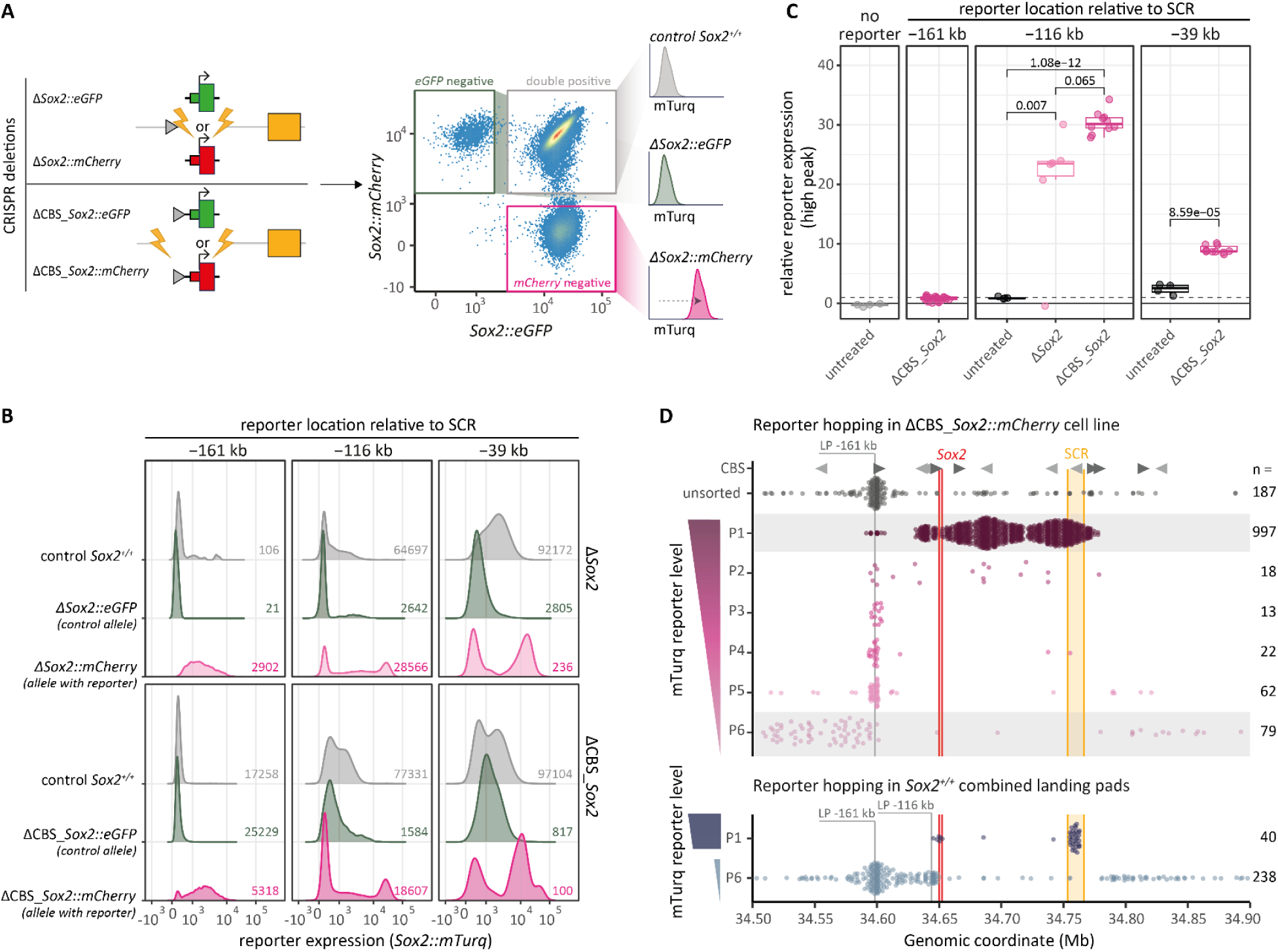
Sox2 gene strongly confines the activation realm of the SCR. **(A)** Left panel: Cas9-mediated deletion of tagged *Sox2* alleles, with or without CBS (grey triangle); right panel: gating and analysis strategy. **(B)** Distribution of mTurq reporter expression levels in three cell lines with reporter inserted in positions −161 kb, −116 kb, and −39 kb, each treated and analyzed as in **A**. Numbers indicate the number of cells measured. **(C)** Maximum relative reporter expression of clonal cell lines after *Sox2::mCherry* or CBS_*Sox2::mCherry* deletion, at reporter locations as indicated. The maximum relative reporter expression is the maximum expression peak indicated in **Fig. S4C-F** (vertical lines), which is then autofluorescence subtracted, normalized to *Sox2::GFP*, and then normalized to the median of the standard −116 kb reporter cell line measured on the same day. Dashed line indicates 1 (standard cell line). Each dot represents one clone (CRISPR edited clones) or one measurement day (untreated clones). **(D)** Top: hopping of reporter (Sox2P) from the −161 kb launch pad in a ΔCBS_*Sox2::mCherry* cell line. Mapped integrations from a large pool of unsorted cells (ctrl) and the six sorted populations P1-P6 (same as in **Fig. 2B**). Data are from 2 biological replicates combined. Grey shading indicates the populations with reporter expression levels that are most different from the starting expression of the unhopped reporter. Bottom: the P1 and P6 sorted integrations combined from hopping in the original −161 kb (2 replicates) and −116 kb (3 replicates) reporter cell lines with *Sox2::mCherry* intact (same data as in **Fig 2D** and **2H**). Grey triangles indicate CBSs and their orientation.

Compared to cells that retained both *Sox2* alleles, cells which only lost *Sox2::mCherry* showed strong upregulation of the reporter (on the same allele) in all three locations (**Fig. 4B**). This upregulation did not occur in cells lacking the *Sox2::eGFP* allele, indicating that the effect is strictly *in cis*. The increase in reporter expression was similar between the Δ*Sox2* and ΔCBS-*Sox2*. The upregulation of the reporter at position −161 kb is remarkable, as it indicates that the endogenous *Sox2* gene normally prevents the SCR from reaching sequences upstream, outside of the TAD. For reporters in the other two positions only a sub-population of cells showed pronounced upregulation (**Fig. 4B**). The basis of this bimodality is unclear.

To gain better understanding of the relationship between reporter location and activation, we sorted out pools and clones of *Sox2::mCherry* negative cells (*cis-*deletion) with high reporter activity. After cell expansion, the reporters at −161 kb and −39 kb displayed stable and uniform reporter expression (**Fig. S4B-D**). In contrast, the −116 kb reporters again exhibited a bimodal expression pattern: a fraction of cells lost reporter expression after sorting the pools (**Fig. S4B**) and clones (**S4E-F**). Comparison of the highest reporter expression peaks of the sorted clones shows that the highest expression was not reached by the reporter closest to the SCR (−39 kb), but rather by the reporter closest to the native location of *Sox2* (−116 kb) (**Fig. 4C**). Overall, these results underscore that the region around the endogenous *Sox2* gene is optimal for high expression, even when this gene including its promoter is deleted.

To further explore the impact of the native *Sox2* gene on the regulatory landscape throughout the locus, we repeated the reporter hopping experiment from the −161 kb launch site in a ΔCBS-*Sox2::mCherry* clone (**Fig. S4C**). Since the founder cell line has intermediate reporter expression (**Fig. S4G**), hopped integrations were expected to be most enriched in the highest (P1) and lowest (P6) expression gates. In the presence of the endogenous *Sox2::mCherry* (*Sox2*^+/+^), P1-level integrations were rare and only found at the SCR and endogenous gene (**Fig. 4D**, bottom). In contrast, when the endogenous *Sox2* gene is deleted, such highly active reporter integrations are found throughout the *Sox2*-SCR range and even extend beyond the cluster of CTCF binding sites ∼7 kb upstream of the deleted *Sox2* promoter. Most silent (P6) reporters are only found at least 55 kb upstream of the deleted gene and 38 kb downstream of the SCR, indicating that the region permissive for activity expanded by roughly 50%, extending beyond the original TAD-boundaries (**Fig. 4D**, top). Together, these data demonstrate that the *Sox2* gene strongly constrains both the level of activation of a competing promoter and the realm of influence of the SCR.

### Unbalanced competition is partially driven by the *Sox2* gene body

While the endogenous *Sox2* gene significantly restricts reporter expression, we found that the reciprocal impact of the reporter on the *Sox2* gene is very minor. First, the reporter inserted in the launch site at −116 kb did not detectably affect *Sox2::mCherry* expression (**Fig. S2A**). Second, among all clonal lines with a hopped reporter, *Sox2::mCherry* expression is remarkably robust. Even in clones with a reporter in the center of the SCR (with ∼10-fold higher expression than at −116 kb) *Sox2::mCherry* expression is reduced by only ∼25% (**Fig. 5A**). This is extremely modest compared to the 3-30 fold upregulation of the reporter upon loss of *Sox2::mCherry* (**Fig. 4C**). Hence, the endogenous gene affects the reporter considerably more than *vice versa*.

**Fig. 5.**
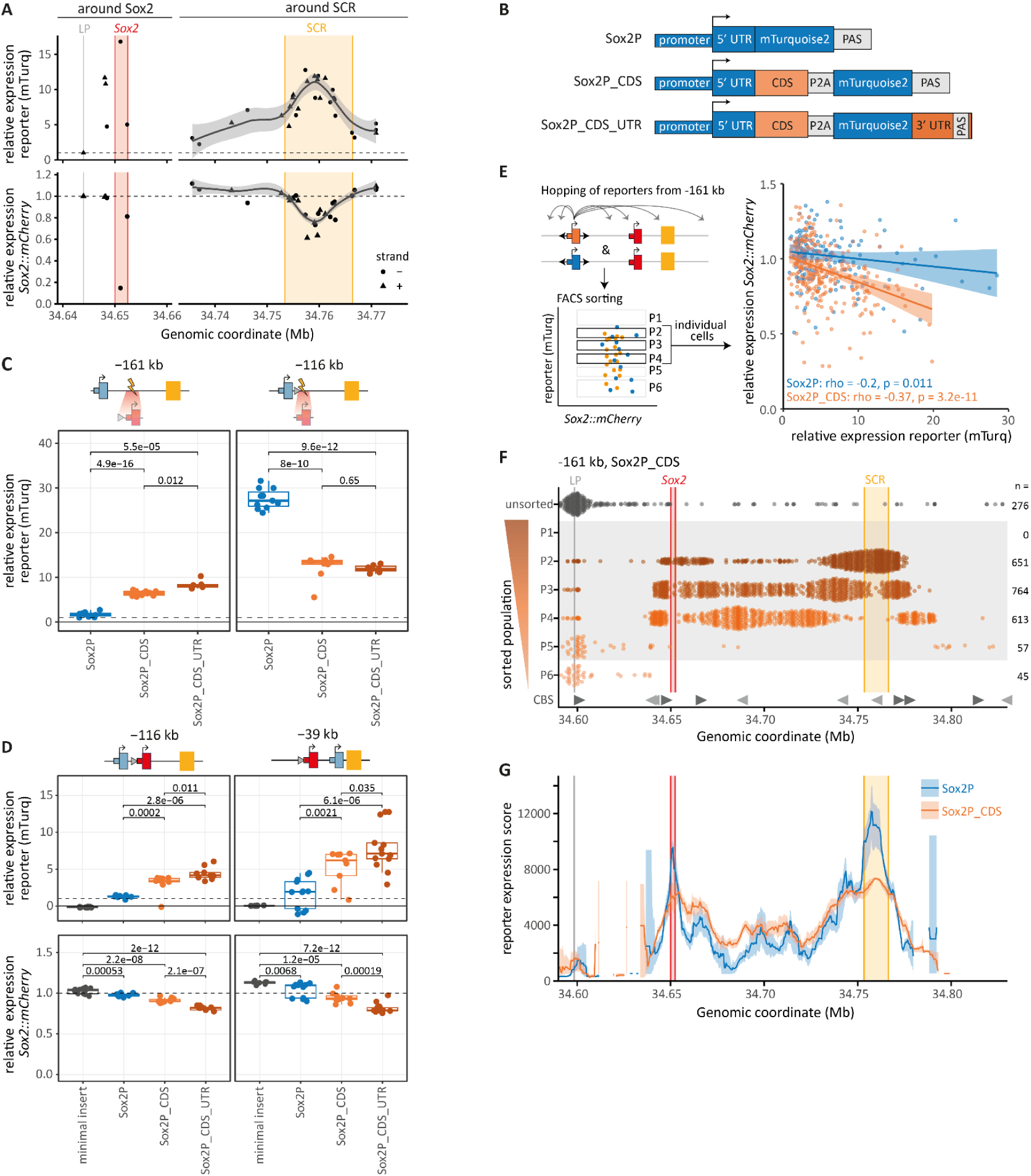
Unbalanced competition is partially driven by Sox2 gene body. **(A)** Relative reporter (mTurq) and *Sox2::mCherry* expression in clones around *Sox2* and the SCR (median expression, estimated autofluorescence subtracted, normalized to GFP, and then normalized to the unhopped reporters at −116 kb). Loess curve shows the trend in expression. Same clones and mTurq data as in **Fig. S2D**, bottom. **(B)** Design of original *Sox2* promoter reporter (Sox2P) and two coding sequence (CDS)-containing reporters. PAS, poly-adenylation signal. **(C)** Relative expression levels of the three reporters at −161 kb and −116 kb, in a ΔCBS_*Sox2::mCherry* cell line. Each dot represents an individual clone. Relative expression was calculated as in **A**. Dashed line indicates 1 (control cell line). **Fig. S5B** shows the same expression values, not normalized to *eGFP*. P-values according to Welch’s unequal variances *t*-test.

We wondered what might give the endogenous *Sox2* gene this competitive edge. The reporter contained the exact promoter and 5’ UTR sequence from the *Sox2* gene, but it lacked the 1.0 kb coding sequence (CDS) and the 1.1 kb 3’ UTR and polyadenylation signal (PAS) (*Sox2* has no introns). Re-analysis of high-resolution region capture micro-C data (*18*) indicates that the SCR not only contacts the promoter of the endogenous *Sox2* gene, but also the gene body (**Fig. S5A**), indicating that the latter sequence might play a role. We therefore generated two new reporters in which we added the *Sox2* CDS alone or in combination with the 3’ UTR/PAS (**Fig. 5B**). We kept the *mTurquoise2* sequence separated from the CDS by a P2A cleavage signal (*24*), so the stability of the fluorescent protein would not be affected by a fusion with the SOX2 protein.

We first tested these new reporters in positions −161 kb and −116 kb after deletion of *Sox2::mCherry*. Strikingly, across multiple independent clonal cell lines, the CDS and CDS/3’UTR additions boosted reporter activity in the −161 kb location by 6-8 fold (**Fig. 5C, Fig. S5B**). Reporter insertion in the −1 16 kb location, just 6 kb upstream of the deleted *Sox2* gene, is a close approximation of replacing the gene with a reporter. Here, the promoter-only reporters showed very high but bimodal expression (**Fig 5C, S5B, S5C**), as observed before (**Fig S4B, E, F**). Surprisingly, the CDS-containing reporters had lower but stable expression (**Fig. 5C, S5C**). We speculate that this lower expression is the result of a negative feedback mechanism on high SOX2 protein levels, which can be triggered by the CDS-containing reporter but not the original *Sox2P* reporter (*25, 26*). Overall, these results suggest that the regulation of the CDS-containing reporter more closely resembles the native *Sox2* gene.

Next, we tested the effect of the CDS when the competing *Sox2::mCherry* gene is also present in the locus. In both the −116 kb and −39 kb position, clonal cell lines with the CDS-containing reporters had 3-7 fold higher expression than those with the original reporter (**Fig. 5D** top, **Fig. S5D**). We then asked whether the presence of the CDS also shifts the competitive balance with the endogenous *Sox2* gene. Indeed, in these cells the addition of the CDS to the reporter caused a 13-18% reduction in *Sox2::mCherry* expression, while additional inclusion of the 3’ UTR in the reporter even caused a 21-31% reduction. (**Fig. 5D** bottom, **Fig. S5D**).

To test whether this competitive characteristic of the CDS reporters is a general feature throughout the locus, we mobilized the CDS-containing reporter from the −161 kb location – where it is not detectably expressed (**Fig. S5E**) – in cells carrying the intact *Sox2::mCherry* (**Fig. 5E**). We compared the relative *Sox2::mCherry* and reporter expression of individual cells with active reporters, as measured by flow cytometry. If the reporter competes with *Sox2::mCherry*, this should be reflected in reduced expression of the latter when the reporter is more active. With the reporter lacking the CDS, this effect is detectable but weak (**Fig. 5E, S5F**). However, when the CDS is present, this effect is about three times stronger (p = 0.008), as indicated by the respective slopes of a fitted regression model (**Fig. 5E, S5G**). Thus, across the random active locations sampled in this hopping experiment, the CDS strengthens the reporter in its competition with the *Sox2* gene.

To further assess if the presence of the CDS in the reporter can change its activation landscape throughout the locus, we mapped the integrations in cells sorted for different reporter expression levels from the CDS hopping experiment (**Fig. S5H**). The sorted integrations of CDS-containing reporter follow a similar pattern as the original reporter (**Fig. 5F**), indicating that the two reporters are stimulated by the SCR in a similar way. However, we did not detect any cells with the highest reporter expression (P1), causing flattening of the activity landscape near the SCR and the endogenous *Sox2* location. This is possibly explained by the same negative feedback that we proposed above. Despite this, the CDS boosted the reporter activity throughout the *Sox2*-SCR range (excluding *Sox2::mCherry* and SCR regions) (**Fig. 5G**). Together, these data indicate that the CDS has two effects on promoter activity: it can boost expression independent of competition; and it increases the competitive edge of a promoter.

## Discussion

Here we developed a novel high-throughput “hopping” approach to create highly detailed quantitative maps of the activation potential of a native genomic locus, based on thousands of reporter integrations. We expect that this approach may also be used to systematically relocate other regulatory elements, such as enhancers, CBSs and other structural elements, and a wide range of synthetic elements.

Earlier probing at low resolution in mouse embryos suggested that promoters respond to an enhancer whenever they are in the same TAD (*11, 27*). Systematic insertion of enhancers and promoters in heterologous genomic loci supported this notion, and pointed to a monotonous decay of promoter activity with increasing genomic distance to the enhancer (*8, 9, 28*). Our high-resolution functional maps of the native *Sox2* locus confirm that the influence realm of the SCR coincides with the TAD, but also reveal an intricate landscape in which pre-existing SCR contact frequency rather than genomic distance is a key determinant of promoter activity. Moreover, the region around the endogenous *Sox2* gene is a “sweet spot” for activation, even in the absence of the gene itself. Possibly, synergistic interactions between the SCR and enhancer-like elements surrounding the gene (*16*) contribute to this.

Another striking result is that the *Sox2* gene is not only passively controlled by the SCR, but also actively constrains the SCR, as indicated by its strong dominance over the reporter. Various studies of native and artificial loci, both in *Drosophila* (*29, 30*) and mammals (*31–33*), showed competition between two promoters for activation by one enhancer. In each case, the enhancer preferentially activated the closest promoter. In contrast, our results show that the SCR strongly favors the native *Sox2* gene regardless of the location of the reporter. Surprisingly, this preference is in part mediated by the *Sox2* coding sequence. Possibly the CDS DNA sequence binds one or more transcription factors that boost SCR interactions; or the transcribed CDS RNA facilitates SCR interactions, as has been proposed for other transcripts (*34*). Our functional maps of SCR – gene interplay lay the foundation for further mechanistic dissection.

## Acknowledgments

We thank the Giorgetti lab for sharing the *Sox2* fosmid; the NKI Genomics, Flow Cytometry, Protein Production and Research High Performance Computing core facilities for excellent support; Marantha Kaagman for valuable experimental contributions; and Elphege Nora, Luca Giorgetti, Elzo de Wit, Tineke Lenstra and members from our lab and the divisions of Gene Regulation and Molecular Genetics for inspiring and helpful discussions.

## Author Contribution

Conceptualization: M.E., C.J.I.M., B.v.S.

Wet-lab experiments: M.E., C.J.I.M., L.D.

Data analysis and visualization: C.J.I.M., M.E.

Method development: M.E., L.D.

Code development: C.L., C.J.I.M.

Writing: M.E., C.J.I.M., B.v.S.

Funding Acquisition: B.v.S., C.J.I.M.

Supervision: B.v.S.

## Data and materials availability

Laboratory notebooks and supplementary data (including primer sequences, gRNAs, plasmids, etc.) are available on Zenodo (10.5281/zenodo.13350269). Raw sequencing data is available on GEO under accession numbers GSE275427.

Code and supplementary files used as input for the scripts are available on Zenodo (10.5281/zenodo.13362255).

All plasmids used will be made available to Addgene.

## MATERIALS AND METHODS

### A. EXPERIMENTAL PROCEDURE

#### Cell culture

F121/9 (CAST/EiJ x S129/Sv) (RRID:CVCL_VC42) female mouse embryonic stem cells (mECSs) F1 hybrid cell line (*35*) and derived clones were cultured in Serum+Lif+2i condition. Briefly, cells were cultured on gelatin-coated (0.1%) culture plates in Glasgow minimum essential medium (Sigma-Aldrich, G5154) supplemented with 15% fetal bovine serum (Thermo Fisher Scientific, 10270-106), 1% L-glutamine (Thermo Fisher Scientific, 25030024), 1% sodium pyruvate (Thermo Fisher Scientific, 11360-70), 1% MEM non-essential amino acids (Thermo Fisher Scientific, 11140-50), 1% penicillin-streptomycin (Thermo Fisher Scientific, 15070063), 100 µM ß-mercaptoethanol, 10 x 4 U leukemia inhibitory factor (LIF; esg1107, Millipore), 1 μM MEK inhibitor PD0325901 (Mirdametinib, MedChemExpress), 3 μM GSK-3β inhibitor CHIR99021 (Laduviglusib, MedChemExpress) in 5% CO_2_ at 37 °C. Cells were passaged every 2 days. Mycoplasma contamination was ruled out by regular testing (#LT07-318; Lonza).

#### Genetic engineering of the *Sox2* locus

##### Tagging endogenous Sox2 alleles

The alleles of the *Sox2* gene were tagged by direct fusion with eGFP and mCherry using CRISPR/Cas9 mediated homologous recombination. We used a previously described plasmid for targeting *Sox2* (*Sox2* sgRNA, Addgene #175553) together with donor plasmids designed to include eGFP or mCherry in between two homology arms (200 bp homologies) for the *Sox2* gene. In brief, 1×10^6^ F121/9 mESCs were transfected with 1 µg of *Sox2* sgRNA and 2 µg of repair template (1 μg of eGFP + 1 µg of mCherry) using 9 µL Lipofectamine2000. Cells were plated in Serum+Lif+2i medium together with a DNAPKcs inhibitor (M3814, MCE, HY-101570) and cultured for two days. Subsequently, mCherry+/eGFP+ cells were sorted sparsely into a 24-well plate (cultured without inhibitor) and resulting colonies were picked and genotyped by PCR. Then they were re-measured by flow cytometry (LSRFortessaTM) using the following lasers and filters: eGFP 488nm BL[B] 530/30, mCherry 561nm YG[D] 610/20. Allele-specific genotyping of *Sox2* was performed by PCR amplification from the fluorophore to the genome across CAST/129S1 SNPs (primers: EMp324, EMp325, MEP68). A clone with the CAST allele *Sox2* gene fused with eGFP and the 129S1 allele *Sox2* gene fused with mCherry was selected as the parental clone.

##### Establishing −116 kb HyTK launch pad

We cloned the vector pME015 containing an mouse *PGK* promoter driving the expression of a double selection marker HyTK (HyTK amplified from plasmid #11684, Addgene), flanked by FRT and F3 heterotypic recombination sites embedded between Sleeping Beauty 5’ and 3’ inverted terminal repeats (ITRs). For establishing the −116 kb launch pad, we used the Alt-R^TM^ CRISPR-Cas9 system from IDT and designed a crRNA that directs cutting 6 kb upstream (−116 kb from SCR) of the endogenous *Sox2*. The crRNA was assembled into an RNP together with the AltR CRISPR tracrRNA (1073189, IDT) and Cas9 nuclease (1081058, IDT). We transfected 50×10^3^ parental cells (*Sox2* GFP+/mCherry+) with the RNP and the PCR product using lipofectamine CRISPRMAX Cas9 Transfection Reagent (CMAX00001, Thermo Fisher Scientific) according to the manufacturer’s protocol. Two days after transfection, Hygromycin was added to the medium (200 μg/mL, 10687010, Invitrogen) to select for HyTK-expressing cells. We picked colonies 6 days later and screened clones for cassette integration by PCR and Sanger sequencing.

##### Establishing −39 kb HyTK launch pad

We created a launch pad in-between *Sox2* and SCR (−39 kb relative to SCR) by CRISPR/Cas9 mediated homology directed repair. For this, we added 1 kb homology arms of the target region upstream and downstream of the HyTK construct (pME015) to generate plasmid pME024, assembled the crRNA with the tracrRNA and Cas9 protein as explained above, and co-transfected the RNP complex together with the pME024 repair template into parental cells (*Sox2* GFP+/mCherry+). Similar as above, cells were selected with Hygromycin and colonies were picked and genotyped by PCR and Sanger sequencing.

#### RMCE donor plasmids

##### Minimal insert

We generated 2000 random 50 nucleotide sequences in silico with a GC content of 50% and scanned them by FIMO for transcription factor binding sites (*36*). We selected one sequence without any transcription factor binding sites, added FRT and F3 sites to the start and end of the sequence and ordered it as an DNA ultramer from IDT. We amplified this sequence by PCR with primers that have 40 bp overlap to the digested ends (BamH1+XhoI) of the target plasmid (pCR Zero, #120275, Addgene) (primers: MEP190, MEP191). Finally we used Gibson assembly to assemble the linearized pCR-Zero vector together with the amplified PCR product according to the manufacturer’s protocol. Two μL of Gibson assembly mix were transformed into 50 μL DH5α (NEB, C2987H) chemically competent cells. Clones were screened by colony PCR and validated by Sanger sequencing.

##### Sox2 reporter construct

We amplified 2346 bp of the *Sox2* promoter (containing the 5’UTR) from a fosmid (Wl1-1017O07, position mm9: chr3:34535173-34570689, kindly provided by Luca Giorgetti’s lab) by PCR using Phusion Polymerase (F-530L, Thermo Fisher Scientific) together with DMSO and GC rich buffer (primers: MEP145, MEP150). The resulting PCR product was cloned into pCR Zero vector (#120275, Addgene) digested with StuI (R0187S, NEB) and transformed into 50 μL DH5α (C2987H, NEB) chemically competent cells. Resulting colonies were picked and genotyped by PCR and validated by Sanger sequencing. To generate the *Sox2P*-mTurquoise2 plasmid, the *Sox2*-promoter-containing pCR Zero plasmid was digested with Xbal + SalI-HF (R3138S, NEB) and ligated (T4 Ligase 5 U/µL, 9015-85-4, Roche) with the Xbal+SalI-HF-digested mTurquoise2-plasmid (#118617, Addgene), resulting in a plasmid containing the *Sox2* promoter with the mTurquoise2 fluorophore. Finally, an SV40 polyA signal was cloned 3’ of the mTurquoise2 (primers: MEP171, MEP172) and F3 and FRT heterotypic Flp-recombinase sites were added by PCR (primers: MEP147, MEP177), resulting into the final pME034 plasmid. The plasmid was validated by Sanger sequencing.

##### Sox2-CDS reporter constructs

We generated two additional *Sox2* reporters containing the *Sox2* coding sequence (957 bp) and either the endogenous *Sox2* 3’UTR or the same polyA signal (SV40) as the original Sox2 reporter construct (pME034). We linearized the reporter plasmid (pME034) with SalI-HF & EcoRV-HF, and PCR amplified the mTurquoise2 gene ± polyA signal from pME034 (primers: MEP224, MEP224, MEP231) and *Sox2*-3’UTR (primers: MEP226, MEP230) and *Sox2*-CDS (primers: MEP222, MEP223) from the same fosmid as before (Wl1-1017O07). A P2A sequence with linker was added by the forward primer for the mTurquoise2 gene (MEP224). We used Gibson assembly to create either the *Sox2*-CDS-polyA(SV40) or the *Sox2*-CDS-3’UTR plasmids. Next, we checked the resulting plasmids by colony PCR and subsequent Sanger sequencing. The final plasmids were named pME040 (F3_*Sox2*_CDS_mTurq_3’UTR_FRT) and pME041 (F3_*Sox2*_CDS_mTurq_polyA_FRT).

#### Recombination-mediated cassette exchange (RMCE)

Inserts were loaded into the SB cassette by recombination-mediated cassette exchange (RMCE). 300,000 cells were transfected with 0.5 μg of Flp recombinase-encoding plasmid (Addgene #13787) and 2 μg of donor plasmid, using 7.5 μL lipofectamine 2000 (Invitrogen, 11668019). Two days after transfection, pools were seeded sparsely and ganciclovir (2.5 μg/mL) was added to the medium to kill HyTK-expressing cells. Resistant colonies were picked, expanded, and screened by PCR. To replace the SB insert in a Sox2P-containing clone derived by SB hopping or CRISPR editing, we first exchanged the Sox2P reporter for the original HyTK cassette by co-transfecting 400,000 to 500,000 cells with 0.5 μg of Flp recombinase-encoding plasmid and 2 μg of a HyTK-containing plasmid (pME015). Two days after transfection, pools of HyTK-expressing cells were selected by adding Hygromycin to the medium. Next, various constructs were loaded into these HyTK-containing pools using RMCE, as explained above.

##### Establishing Sox2-CDS clones

Clones containing the Sox2P, Sox2P-CDS or Sox2P-CDS-UTR construct in different launch pads (−39 kb, −116 kb, −161 kb) were generated by RMCE and single cell sorted in 96 well plates. Subsequently, clones were genotyped by PCR. Only clones containing the right insert and genomic background were measured by flow cytometry (*Sox2::mCherry, Sox2::eGFP*, reporters) and used for downstream analysis.

#### Establishing proof of principle cell line

For a different project we used a plasmid containing a stretch of 200 random nucleotides (free of any CTCF motifs), followed by a CTCF binding site flanked by LoxP sites (kindly provided by Luca Braccioli, Elzo de Wit group). We PCR amplified the insert with flanking F3/FRT sites (primers: MEP74, MEP75) and cloned the resulting PCR product into StuI blunt cut pCR-Zero vector, creating donor plasmid pME013. Next, we used RMCE to integrate the insert into the −116 kb launch pad of *Sox2* GFP+/mCherry+ F121/9 mESCs, resulting in a cell line called CBS16. To generate a ΔCTCF control cell line (CRE6), we transfected the cells with a CRE-Puro plasmid (pJK14-946-CRE-Puro), selected with Puromycin and seeded surviving cells as single cells in 96 well plates by FACS (BD FACSAriaTM Fusion Flow Cytometer). Resulting clones had an insert of 282 bp in between FRT/F3 sites and were genotyped for the floxed CTCF motif by PCR and validated by Sanger sequencing. Those cells were used for the proof of principle experiment. The sequence of the resulting insert (200 arbitrary nucleotides and one loxP site) can be found in the supplementary data sheet.

#### SB Hopping

##### SB hopped cell pools

For each replicate and cell line, SB hopping was induced by transfecting 2.5 to 5 million cells with the bicistronic plasmid (pME07) encoding SB transposase SB100x and the human nerve growth factor receptor (LNGFR), using 15 µL Lipofectamine 2000 (Invitrogen, 11668019) and 5 µg of plasmid per million cells. As a control, 0.5 million cells were transfected with a similar bicistronic plasmid encoding PiggyBac (PB) transposase and LNGFR (pLD042). After 30 hours, transfected cells were enriched using MACS MS columns (130-042-201, Miltenyi Biotec) and LNGFR MicroBeads (130-091-330, Miltenyi Biotec) according to the manufacturer’s protocol. One week after transfection, 0.2 million SB-transposase-transfected cells were set apart as an unsorted pool and the remaining cells were sorted by FACS (BD FACSAriaTM Fusion Flow Cytometer) into pools for populations based on mTurq expression level, using the following lasers and filters: mTurq: 442nm V[F] 470/20, eGFP: 488nm BL[B] 530/30, mCherry: 561nm YG[D] 610/20 (see Fig 2B for the exact gates). PB transposase transfected cells served as mock-treated FACS control. In the first biological replicate of the −116 kb Sox2P-reporter (**Fig 2B, 3A**) cells were only selected based on mTurq level, in all later replicates we selected only *Sox2::mCherry*-positive cells for all *Sox2::mCherry* containing cell lines and only *Sox2::mCherry*-negative cells for the ΔCBS_*Sox2::mCherry* cell line. Cells were sorted into up to 5 pools of up to 1000 cells per pool. Sorted pools were expanded and crude lysates were obtained from a full 96 well plate by lysing with 50 µL DirectPCR Lysis Reagent (102-T, Viagen Biotec) supplemented with 100 µg/mL proteinase K, and incubating 2.5 h at 55 °C and 45 min at 85 °C. After expansion, gDNA was extracted from the unsorted control cell pools using the ISOLATE II Genomic DNA Kit (BIO-52067, Bioline).

##### Establishing clones

To generate a panel of clonal cell lines with active reporters, we FACS sorted single cells either from previously sorted P1 and P2 pools (biological replicate 1) or directly from the full pool of SB-transposase-transfected cells (replicates 2 and 3, only deletion clones). mCherry-positive and mCherry-negative cells were sorted to obtain clones with hopped insertions and hopping-induced deletions, respectively. When sorting from the full pool of SB-transfected cells we also gated for high (P1) mTurq expression. The cell line with a reporter integration at −161 kb was isolated in an earlier SB hopping experiment (not described here), by sorting out mTurq-negative cells from a pool of SB-transposase-transfected cells (from the −116 kb reporter cell line). Sorted clones were expanded and crude lysates were obtained from a full 96 well plate, as for the cell pools. Deletion clones presumably derived from one hopping event (from the same experiment with identical location and orientation) were averaged.

##### Genotyping hopping-induced deletion

A few hopping-induced deletion clones were genotyped to confirm the genomic deletions. We used PCR (primers: CMP116-121) to amplify three regions containing annotated SNPs on the CAST allele and Sanger sequencing to evaluate which allelic fragment(s) got amplified. By examining the Sanger sequencing traces, we could categorize if and which allele got deleted (**Fig. S3B**).

#### Tagmentation mapping

SB integrations were mapped by amplifying and sequencing the ITR-genome junction using a tagmentation-based approach, based on (*19*). The procedure was similar to (*37*), but using the following SB-ITR specific primers: MEP009 (5′ ITR, reverse) or LD027 (3′ ITR, forward) for the linear enrichment and MEP011 (5′ ITR, reverse) or MEP034 (3′ ITR, forward) for PCR1. Input for one tagmentation reaction was either 100ng gDNA or 3 µL crude lysate. The 72 °C extension times of all PCRs were increased to 1 min. In addition, the composition of the linear enrichment PCR was changed to 10 μL of tagmented DNA, 2 μL of 1 μM primer, 4 μL dNTPs (10mM), 8 μL 5x Phusion® HF Buffer (NEB), 0.5 μL Phusion® HS Flex polymerase (2 U/μL - NEB), in a final volume of 40 μL. For all clones and some experiments with sorted pools (proof-of-principle experiment, third replicate of the reporter hopping) we halved this reaction to a final volume of 20 µL. For the unsorted control pools, 10 separate tagmentation reactions and library preparations were performed, usually using one set of sequencing indices. Libraries were sequenced with 150 bp paired-end reads using either an Illumina MiSeq system (reagent kit v2) or Illumina NextSeq 550 system (mid-output kit) including 10% of PhiX spike-in.

#### CRISPR deletions

##### Deletion endogenous Sox2

To delete the endogenous *Sox2* gene, we designed two gRNAs targeting upstream of the CTCF site, one at the start of the *Sox2* promoter (not targeting the reporter) and three downstream of the *Sox2* gene (one in the 3’ UTR, two after). gRNAs were cloned into an expression plasmid containing mCherry. 0.5 million cells were transfected with 1 µg Cas9 plasmid (PX330) and 1.5 µg gRNA plasmid pool (equimolar pool) using 7.5 µL lipofectamine 2000. Transfected cells were isolated after one day, by sorting cells with increased mCherry expression. More than three weeks later, *Sox2* and reporter expression were measured in these cells by flow cytometry (BD FACSAria™ Fusion) and *Sox2::mCherry* negative cells with high reporter expression were sorted as pools. *Sox2::mCherry* negative, reporter-high clones were either isolated at the same time (−116 kb ΔCBS_*Sox2*), or isolated later from the sorted pools by single cell dilution (−116 kb *ΔSox2*) or by FACS sorting *Sox2::mCherry*-negative mTurq-positive cells (−161 kb ΔCBS_*Sox2*, −39 kb ΔCBS_*Sox2*). We isolated gDNA or crude lysates (as before) for all clones and performed a PCR across the intended deletion (*ΔSox2*: CMP075+CMP081; ΔCBS_*Sox2*: CMP077+CMP081). In addition, we performed a PCR from the intact GFP or 3’ UTR to the downstream region (CMP024+CMP081 or CMP080+CMP081) to confirm that the *Sox2::GFP* allele was not inverted. Clones where either PCR failed were excluded from all analyses.

##### SCR deletion

We cloned two previously published gRNA sequences (*13*) targeting 5’ or 3’ of the SCR into a Cas9 expressing px330 backbone (#158973, Addgene) according to the protocol published by the Zhang lab (*38*). Used gRNA sequences were reported to work in mESCs in an earlier publication (*13*). Resulting plasmids were verified by Sanger sequencing and transfected into *Sox2* GFP+/mCherry+ F121/9 mESCs. In brief, 0.5 million cells were transfected with 0.5 µg 5’SCR-sgRNA plasmid and 0.5 µg 3’SCR-sgRNA plasmid using Lipofectamine2000. Cells were analyzed for eGFP and mCherry expression by FACS (LSRFortessaTM) after 5 days of culturing using the following lasers and filters: eGFP BL[B] 530/30, mCherry YG[D] 610/20.

#### Flowcytometry

Allele-specific *Sox2* expression and reporter expression of clones and pools were measured on a BD LSRFortessa™, using the BD™ High Throughput Sampler (HTS) to directly sample from 96 well plates. The following lasers and filters were used: eGFP 488nm BL[B] 530/30, mCherry 561nm YG[D] 610/20, mTurquoise2 405nm V[F] 470/20. Single cells were gated using FlowJo and exported as fcs files which were loaded in R using the flowCore package (*39*) for further analysis.

#### CTCF ChIP

F121/9 (CAST/EiJ x S129/Sv) *Sox2* mCherry+/eGFP+ were cultured in Serum+Lif+2i condition. For chromatin preparation, 10 million cells (per condition) were cross-linked with a final concentration of 1% formaldehyde for 10 min. The cross-linking reaction was quenched with 2.0 M glycine (0.2 M final concentration). The cross-linked cells were then lysed and sonicated to obtain approximately 300 bp chromatin fragments with a Bioruptor Plus sonication device (Diagenode). For ChIP assays, antibodies were first coupled with Protein G dynabeads (10003D, Thermo Fisher Scientific), and the sonicated chromatin was then incubated overnight at 4 °C. After incubation, the beads were washed, the captured chromatin was eluted and cross-linking was reversed. The released DNA fragments were purified with the MinElute PCR Purification kit (28004, Qiagen). The ChIP experiment was performed with the following antibody: CTCF (07-729, Merck Millipore; 5 µL per ChIP). The purified DNA fragments were prepared according to the protocol of the KAPA LTP Library Preparation kit (KR0453, Roche) before sequencing. All ChIP-seq libraries were sequenced with single-end 150 bp cycle mode on an Illumina NextSeq 550 (mid-output kit).

### B.#DATA PROCESSING AND COMPUTATIONAL ANALYSIS

#### Computational analysis of SB hopping

##### Tagmentation mapping

We used the pipeline published in (*12*) to map SB integrations. For experiments using the −161 kb launch pad we mapped to the mouse reference genome (Release M23 – mm10), for the −116 kb launch pad we mapped to mm10 with an *in silico* insertion of the SB cassette (including exact flanking indels from the CRISPR insertion) and lifted over the integration positions to the mm10 genome. In the pilot experiment (**Fig. 2C**), the pipeline was also used to assign integrations to the 129S1 or CAST allele based on SNPs in the mapping reads, using SNP annotation of CAST and 129S1 alleles from the Mouse Genomes Project (https://www.mousegenomes.org/publications/ (*40, 41*)). Reads were filtered for a minimal mapping quality of 10. For the sorted pools, integrations with at least two unique mapping reads were retained. To ensure that high reporter expression is due to location and not due to hopping-induced *Sox2::mCherry* deletions, the integrations in the in the P1 pools were further filtered for having at least one read mapping each from the 5′ ITR and 3’ ITR (reverse and forward read). For the large unsorted control pools we filtered integrations less stringently, including any integration with at least one mapping read. Note that the integrations in these large unsorted pools are not mapped exhaustively. In a few experiments, integrations exactly matching the −39 kb launch site (chr3:34721183-34721192) or an earlier derived hopped clone (chr3:25018987 and chr3:35232946) were found across many pools. This indicated a PCR contamination from an earlier experiment, therefore we filtered out all integrations exactly matching these coordinates.

##### Clones

To identify the true integrations in clonal cell lines, we filtered more stringently, requiring at least ten unique mapping reads. Next, we discarded any locations that were supported by <20% of the read count of the top location, since these are likely to be either index-swapped reads from other clones or off-target reads. To call an insertion with confidence, we required to find exactly one integration supported by 5’ ITR and 3’ ITR mapping. For confident deletion clones, we required to find exactly two locations: one supported by the 5’ ITR mapping, and one by the 3’ ITR mapping. Any clones with more mapped locations were manually reviewed: any clones for which the extra reads could be explained by index-swapping were included, while clones with evidence of a secondary integration or an unexplained high number of mapping reads from a secondary location were discarded.

##### Expression score from sorted pools

To calculate expression scores from sorted pools, we first combined the integrations from each experiment per sorted population. Identical integrations found in multiple sorted pools from one experiment most likely originated from a single hopping event. Still, we treated them as separate data points because they were independently sorted into an expression gate. To calculate the expression score for any cell line, we first took the median fluorescence intensity (Fluo_P_) of each sorted population P1 through P6 that was measured during the FACS sorting process. For −161 kb Sox2P P1 cells were sorted but no cells were recorded (frequency was less than 1 in 100.000 cells), so the mean Fluo_P_ from −116 kb Sox2P P1 was used. Next, for any given window (W), the expression score was defined as the average of these population fluorescence values, weighted by the fraction of all integrations of each population falling within that window 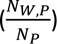 :

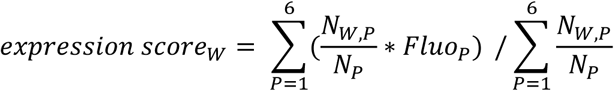

Whenever replicate data were pooled, the median fluorescence per gate varied slightly between biological replicates so the fluorescence of each experiment (E) was weighted by the number of integrations from that experiment:

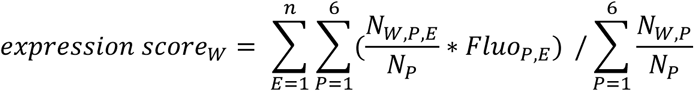

To obtain a running window expression score (e.g. 10 kb window), integrations were counted in smaller bins (e.g. 1 kb, this is the step size) and the expression score was calculated from a group of bins (e.g. 10 bins). Window and step size were selected based on the density of integrations in each experiment and region of interest. To estimate a 95% confidence interval, the genome-wide list of integrations for each cell line was bootstrapped 5000 times and the expression score was calculated from each bootstrap. Reported are the median of these bootstrapped expression scores as well as the 2.5^th^ to 97.5^th^ percentile (95% confidence interval), for each window with at least 3 sorted integrations. To obtain the strand-specific expression score, only integrations on a specific strand were included in integration count of a window (*N_W,P_*), while integrations on both strands were included in the total number of integrations of the population (*N_P_*).

#### Computational analysis flowcytometry

##### FACS color density plots

To show the exact FACS gating used to sort SB-transposase-transfected cells, we loaded the FACSDiva experiment file (.xml) and the raw fcs files in R using diva_to_gatingset from the CytoML package (*42*). Color density plots were created using ggcyto and geom_hex. The other color density plots (e.g. showing *Sox2::mCherry* and *Sox2::GFP* negative populations after CRISPR treatment) were created from gated single cells exported from FlowJo. The threshold for GFP or mCherry negative cells was determined by the 99th percentile of the WT control, while the threshold for GFP or mCherry positive cells was 1.5 times this value.

##### Normalization pools and clones

To compare the *Sox2* and reporter expression levels between clones (**Fig. 3C**), the fluorescent values were corrected and normalized. First, the median fluorescence intensity (MFI) was calculated for each fluorophore and each clone. Next, the MFIs of a mESC control cell line lacking any fluorescent protein were subtracted from the respective MFIs of the other samples, to correct for autofluorescence. Subsequently, the corrected mCherry and mTurquoise2 values were normalized by the corrected eGFP value of the same sample, to adjust for any global effects on *Sox2* expression. Because there was variation in the absolute MFIs between measurement days, we finally divided the normalized fluorescence values by the respective value of the original −116kb_Sox2P cell line, measured on the same day. In this way, we obtained relative *Sox2::mCherry* and reporter (mTurquoise2) expression levels that could be compared between measurement days and experiments (**Fig. 3C, 4C, 5A, 5C-D**). To compare the expression of all fluorophores without adjusting for global changes in SOX2 level, we divided the autofluorescence-corrected MFI values by those of the −116kb_Sox2P cell line (‘relative expression, not normalized’, **Fig. S5B, S5D**). The measurements of the clones derived by SB hopping (insertions and deletions, e.g. **Fig. 3C**) contained no wild-type control, so we used the wild-type control from a different day with closely matching fluorescent values of the original −116kb_Sox2P cell line.

##### Normalization single cells

To compute the relative reporter and *Sox2::mCherry* expression of individual SB-transposase-transfected cells (**Fig. 5E**), the MFIs of the wild-type mESC sample were subtracted from the fluorescent values of each cell. Next, the corrected mCherry and mTurquoise2 values were normalized by the eGFP value of the same cell. Finally, all individual cell measures were scaled to the corrected, normalized MFIs of the −116kb Sox2P cell line to be able to combine the data from two measurement days (biological replicates). Only cells in P2-P4 were used because P5 and P6 contain primarily many non-mobilized reporters (so individual cells do not represent a unique integration location) and no cells were recorded in P1.

##### Bimodal expression patterns

To determine the modes of the fluorescence density distribution in clones with a bimodal reporter expression pattern, we used the function getPeaks from the flowDensity package (*43*) on the bi-exponentially transformed data (as in the figures). We extracted at most two highest peaks and selected the peak with the highest expression value. This mTurquoise2 value was corrected and normalized to compare between experiments as previously explained for MFI values.

#### Computational analysis CTCF ChIP

Raw sequencing data were filtered (>=Q15) and adapters were trimmed using fastP (*44*). Sequences were mapped to mm10 reference genome using the BWA alignment tool (0.7.17-r1188) after indexing the mm10 reference genome. The coverage files (bigwig files) were generated with RPKM normalization using deepTools (version 3.0). Peak calling was performed with MACS2 at a q-value cutoff of 0.05 and CTCF motifs were called using Jaspar (CTCF:MA0139.1) database running MotifScan (v1.3.0). By manual inspection of the CTCF bigwig files we could see that one site was clearly bound but not annotated by the MotifScan. Therefore we added it manually (mm10; chr3: 34772210).

#### External data

#### Region capture micro C

Processed region capture micro-C (RCMC) data from (*18*), aligned to the mm39 genome, was visualized using the GENOVA package (*45*). For the contact matrices in **Fig. 1B** and **Fig. S5A**, all annotations of the *Sox2* locus were lifted over to the mm39 genome. Insulation score was calculated using the function insulation_score, on the 1 kb resolution data, with window = 20 (20 kb window size). Trans-interactions between the *Sox2* gene and SCR were visualized using the function trans_matrixplot, using the 200 bp resolution RCMC data. To correlate contact probability to expression score, we created a virtual 4C on the mm39 genome using as a viewpoint the region containing the two essential enhancer elements in the SCR (mm39: chr3:34811722-34816213, (*16*)) at 1 kb resolution. Then we lifted over the contact probabilities to the mm10 genome build and calculated the expression score on the same 1 kb bins. Only bins with at least 3 reporter integrations were included in the correlation.

## Supplementary Figures

**Fig. S1.**
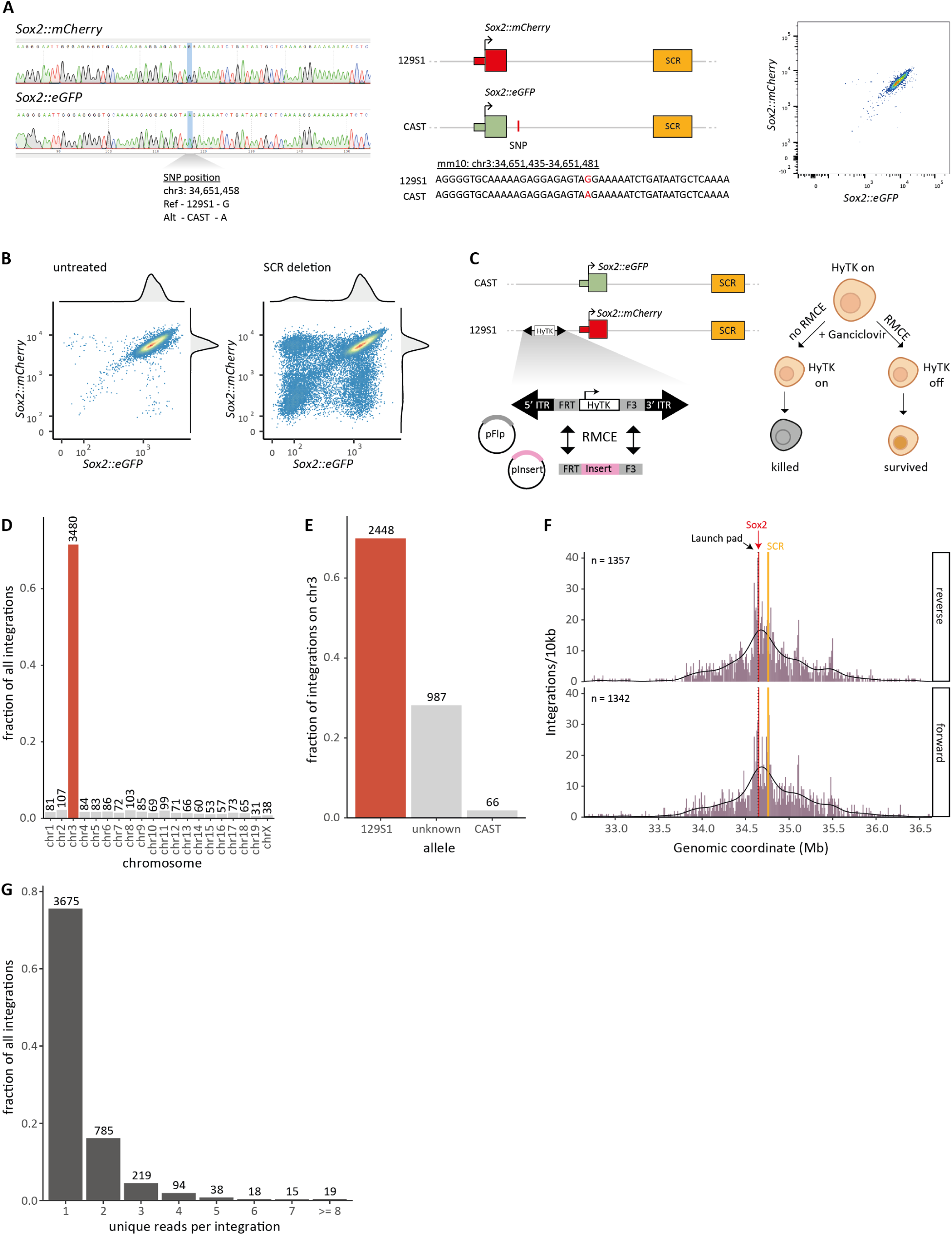
Cell line generation and proof of principle experiment. **(A)** Left and middle: PCR and Sanger sequencing with mCherry and eGFP specific primers indicates on which allele each fluorophore is inserted. Right: expression level of Sox2::mCherry and Sox2::eGFP measured by flow cytometry. **(B)** Allele-specific Sox2 expression in the dual tagged cell line, in cells treated with CRISPR to delete the SCR and in untreated control cells. **(C)** Recombination-mediated cassette exchange: The hygromycin phosphotransferase-thymidine kinase fusion gene (HyTK) is flanked by heterotypic Flp recombinase recognition sites (FRT & F3) and embedded into a Sleeping Beauty transposon (SB-ITRs), integrated via CRISPR-Cas9-mediated knock-in 6 kb upstream of the Sox2::mCherry fusion gene. Co-transfection of the Flp recombinase expression plasmid (pFlp) and donor plasmid (pInsert) with subsequent ganciclovir selection for 7 days results in cells with the intended insert. **(D-G)** Statistics from the proof-of-principle hopping experiment, where a SB transposon carrying a 282 bp arbitrary DNA sequence was relocated. Number of integrations is indicated (above the bars or n). **(D)** Fraction of integrations mapped per chromosome. The launch pad is on chr 3. **(E)** Fraction of all integrations on chr 3 mapped to each allele, or not assigned to either allele. The launch pad is on the 129S1 allele. **(F)** Number of integrations per 10 kb bin, split by SB orientation. **(G)** Distribution of the number of unique mapping reads supporting each integration. **(B, D-G)** Data from one (proof-of-principle) biological replicate.

**Fig. S2.**
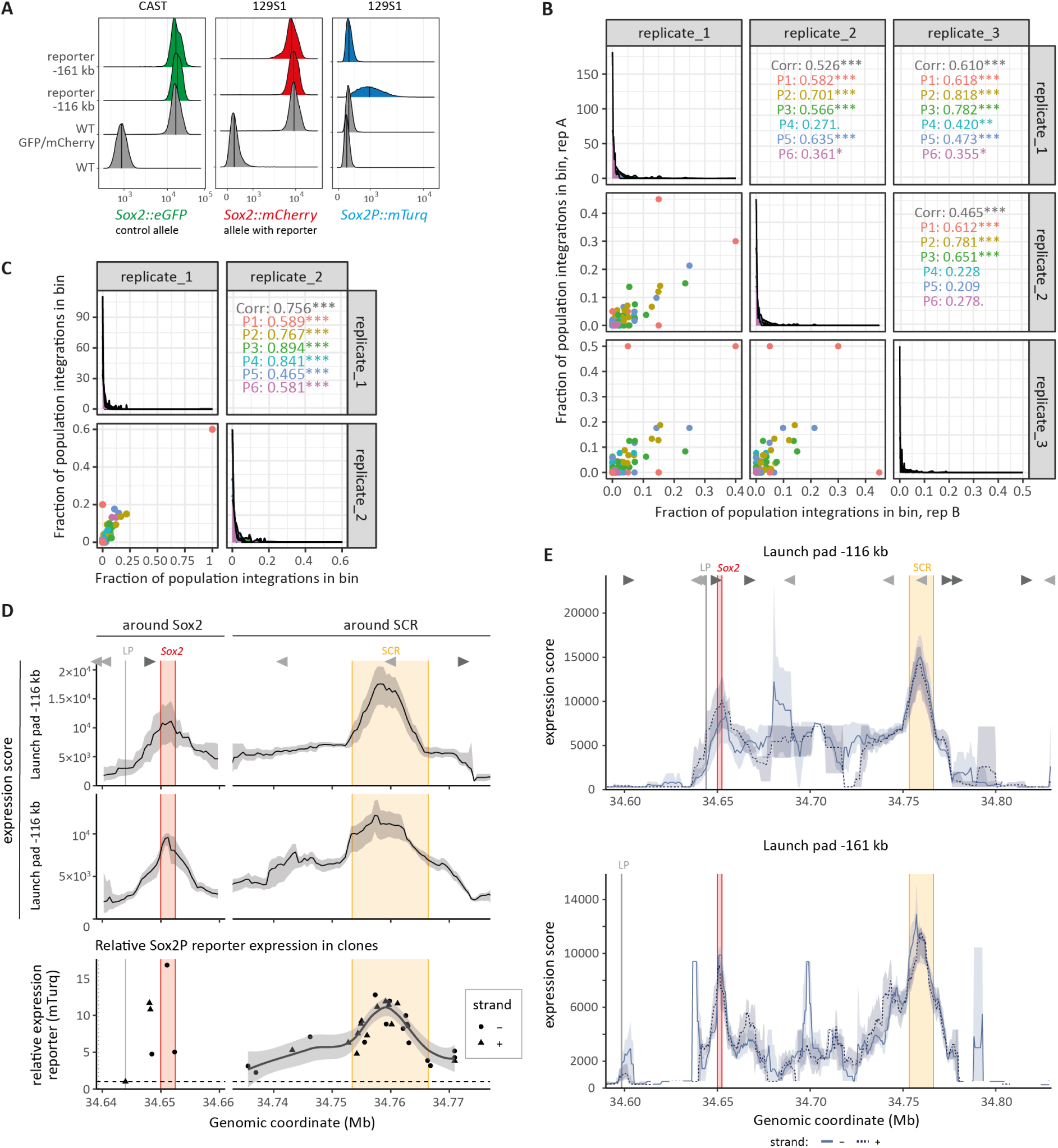
Reproducibility of reporter hopping and validation of the expression score. **(A)** Distribution of expression of *Sox2::eGFP* (green), *Sox2::mCherry* (red) and the reporter (mTurq, blue), in the −161 kb and −116 kb reporter cell lines and the parental control cell lines measured by flow cytometry. **(B)** Reproducibility of the fraction of sorted integrations across the *Sox2* locus, from the −116 kb launch pad. Bottom left panels: each dot represents the fraction of all integrations of that population found in a 5 kb bin in the *Sox2* locus, compared between two biological replicates. Colors indicate the 6 sorted populations. Panels along diagonal: distribution of observed fractions in each replicate. Top right panels: Spearman correlation between two replicates. **(C)** As B, but for the −161 kb launch pad, with two biological replicates. **(D)** Top: the computed expression scores around *Sox2* gene and SCR based on the sorted integrations from the −116 kb and −161 kb launch pad (see **Fig. 2D, H**), smoothened using a 5 kb running

**Fig. S3.**
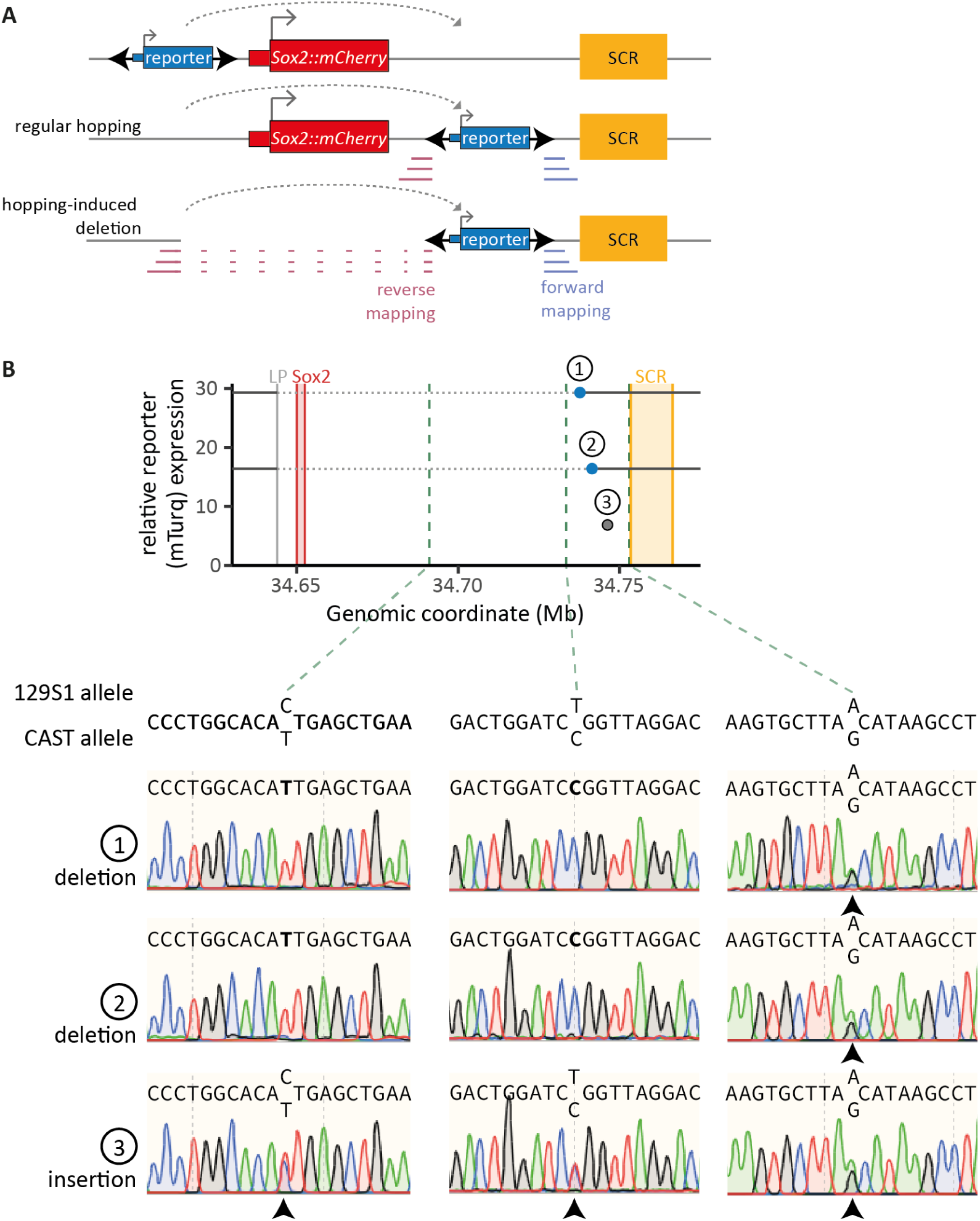
Mapping of hopping induced deletions and insertions. **(A)** Schematic of Tn5-based mapping in cells with a regular hopping event or a hopping-induced deletion. In case of a hopping-induced deletion, one SB ITR will be mapped to the launch site while the other is mapped to the insertion location. **(B)** Confirmation of the loss of the 129S1 allele in the deleted region of deletion clones. Top: genomic location and expression of two hopping-induced deletion clones (1, 2, blue) and one regular insertion clone (3, grey). Horizontal dashed grey lines indicate expected deletions, vertical dashed green lines indicate the locations of three known SNPs between the 129S1 and CAST alleles. Bottom: Sanger sequencing across the three SNPs in the three cell lines. Black arrowheads indicate double peaks at the SNPs, indicating that both alleles are present. When only the CAST allele is present, the SNP is indicated in bold in the sequence above the Sanger trace.

**Fig. S4.**
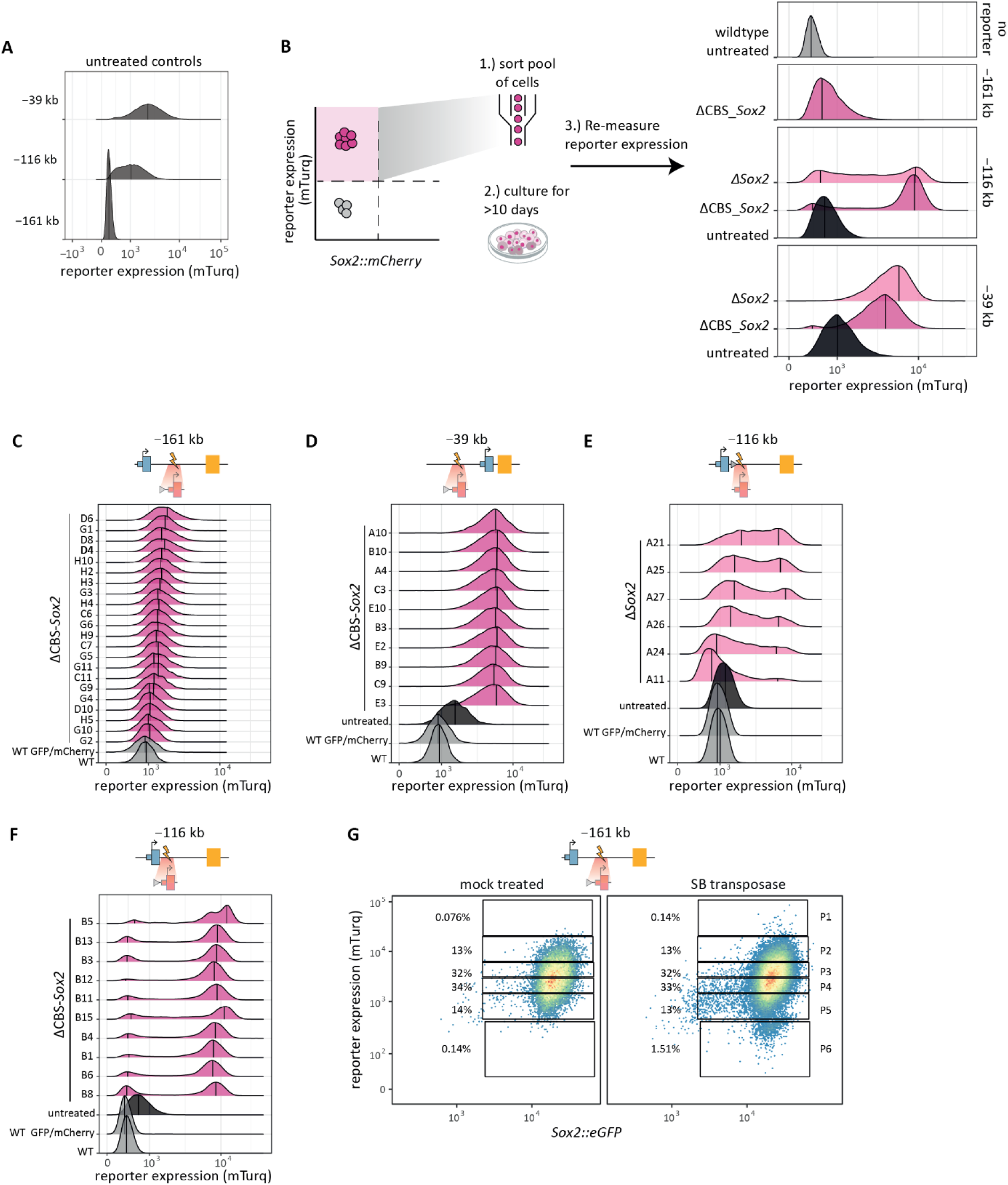
Sox2P-reporter expression in Sox2::mCherry deleted pools and clones. **(A)** Reporter expression in untreated control cell lines (−39 kb, −116 kb, −161 kb). **(B)** Distribution of reporter expression in *Sox2::mCherry*-negative, reporter-high sorted pools. **(C)** Distribution of reporter expression in CBS_*Sox2::mCherry* deleted, reporter-high sorted clones from the −161 kb reporter cell line. Clone D4 (in bold) was used for hopping (see **Fig. 4D**). **(D)** As in C, but for the −39 kb reporter cell line. Untreated reporter cell line is indicated in black. **(E)** As D, but for *Sox2::mCherry* deletion in the −116 kb reporter cell line. **(F)** As D, but for CBS_*Sox2::mCherry* deletion in the −116 kb reporter cell line. **(C-F)** Vertical lines indicate the (at most) two highest peaks in the density distribution, the peak with the highest

**Fig. S5.**
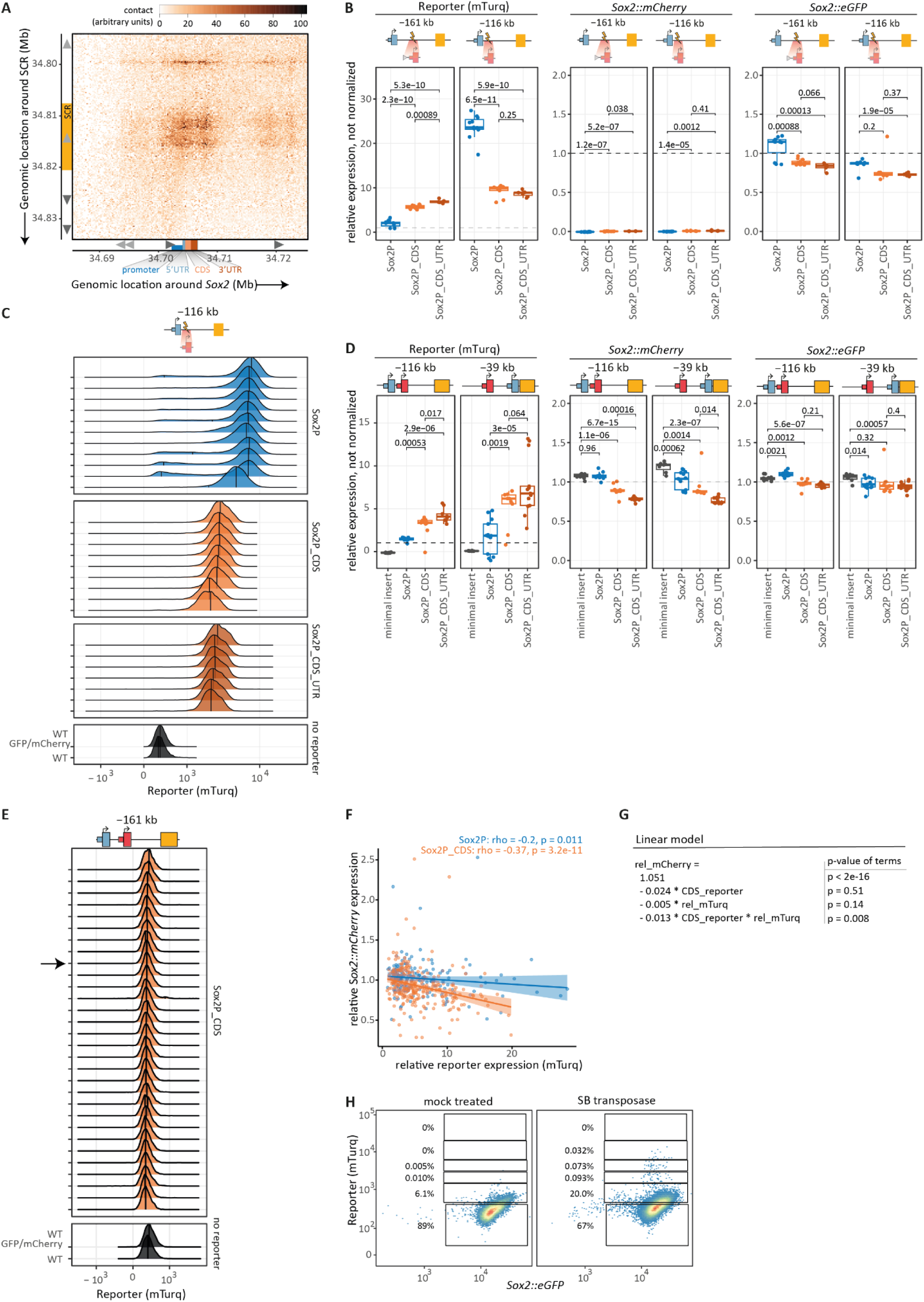
Impact of the Sox2 coding sequence on reporter and *Sox2* expression. **(A)** Contact between *Sox2* and the SCR (region-capture micro-C data [(*18*)], 200-bp resolution). Positions of the components of the *Sox2* gene (blue, orange, red; x-axis) and SCR (yellow; y-axis) are indicated. Grey triangles indicate CTCF binding sites and their orientation. Arrows indicates the genomic direction. **(B)** Relative expression of the reporter (mTurq), *Sox2::mCherry* and *Sox2::eGFP*, not normalized to *Sox2::eGFP*, for the three reporters inserted at −161 kb and −116 kb. Each dot represents one clone. Expression values are autofluorescence subtracted, and then normalized to the control reporter cell line (*Sox2* reporter at −116 kb) measured on the same day. Dashed line indicates 1 (level of the control cell line). P-values according to Welch’s unequal variances *t*-test. **(C)** Distribution of reporter expression of reporters integrated at −116 kb in a ΔCBS_*Sox2::mCherry* cell line. Each density plot is one clone, vertical lines indicate the (at most) two highest peaks in the density distribution. **(D)** As **B**, for the three reporters and a minimal insert (not containing any promoter or mTurq) inserted at −116 kb and −39 kb launch pads with *Sox2::mCherry* intact. **(E)** Reporter expression in a panel of clones with the Sox2P_CDS reporter integrated at the −161 kb launch pad with *Sox2::mCherry* intact. The arrow indicates the clone used for hopping (see **Fig. 5F**). **(F)** Correlation between relative *Sox2::mCherry* and reporter (mTurq) expression from the Sox2P and Sox2P_CDS reporter, from cells with reporter expression in the gates P2-P4 after induction of hopping. As in **Fig. 5E**, but also showing the outlier points. **(G)** Linear model of the data in **F**, describing how the relative *Sox2::mCherry* expression (rel_mCherry) depends on the identity of the reporter (CDS_reporter, is 0 for the Sox2P reporter and 1 for the Sox2P_CDS reporter), and the relative reporter expression (rel_mTurq). Right side indicates the p-values of the coefficients of the model, testing whether they are significantly different from 0. **(H)** Expression of *Sox2::eGFP* (control) and the *Sox2* reporter (mTurq) in the untreated cells and cells transfected with the SB transposase plasmid, for the −161 kb Sox2P_CDS reporter cell line. The percentage of cells in each sorted gate is indicated. Cells are gated for *Sox2::mCherry*-positive single cells. Representative image of 2 biological replicates.

